# Vitamin C activates young LINE-1 elements in mouse embryonic stem cells via H3K9me3 demethylation

**DOI:** 10.1101/2023.08.07.552254

**Authors:** Kevin C.L. Cheng, Jennifer M. Frost, Francisco J. Sánchez-Luque, Marta García-Canãdas, Darren Taylor, Wan R. Yang, Branavy Irayanar, Swetha Sampath, Hemalvi Patani, Karl Agger, Kristian Helin, Gabriella Ficz, Kathleen H. Burns, Adam Ewing, José L. García-Pérez, Miguel R. Branco

## Abstract

Vitamin C (vitC) enhances the activity of 2-oxoglutarate-dependent dioxygenases, including TET enzymes, which catalyse DNA demethylation, and Jumonji-domain histone demethylases. The epigenetic remodelling promoted by vitC improves the efficiency of induced pluripotent stem cell derivation, and is required to attain a ground-state of pluripotency in embryonic stem cells (ESCs) that closely mimics the inner cell mass of the early blastocyst. However, genome-wide DNA and histone demethylation can lead to upregulation of transposable elements (TEs), and it is not known how vitC addition in culture media affects TE expression in pluripotent stem cells. Here we show that vitC increases the expression of evolutionarily young LINE-1 (L1) elements in mouse ESCs. We find that TET activity is dispensable for these effects, and that instead L1 upregulation occurs largely as a result of H3K9me3 loss mediated by KDM4A/C histone demethylases. Despite increased L1 levels, we did not detect increased somatic insertion rates in vitC-treated cells. Notably, treatment of human ESCs with vitC also increases L1 protein levels, which could impact the genetic and epigenetic stability of human pluripotent stem cells.

## Background

A key feature of epigenetic regulation is its susceptibility to environmental inputs. One important mechanism of environmental influence on epigenetics is through the availability of cofactors for modifying enzymes. For example, DNA and histone methylation require S-adenosylmethionine (SAM); histone acetylation and de-acetylation depend on acetyl-CoA and oxidised nicotinamide adenine dinucleotide (NAD+), respectively (1,2). TET methylcytosine dioxygenases, which oxidise 5-methylcytosine (5mC) in a pathway for DNA demethylation, and the Jumonji family of histone demethylases, are dependent on alpha-ketoglutarate/2-oxoglutarate (2OG), oxygen and ferrous iron (Fe(II)) (1,2). These co-factors are therefore powerful modulators of epigenetic effects on cell function and identity. For example, a high 2OG-to-succinate ratio in mouse embryonic stem cell (mESC) culture promotes DNA and histone demethylation, and maintains pluripotency (3). In tumours, hypoxia results in a loss of TET activity, leading to promoter hypermethylation and increasing tumour growth (4).

Vitamin C (vitC) is an additional key modulator of 2OG-dependent dioxygenase (2OGX) activity, which acts through recycling of Fe(III) to Fe(II), and as such can also influence cellular epigenetics through regulation of both DNA and histone demethylation (5–7). The presence of vitC in culture media promotes the pluripotent state in both mouse and human cells (8). Whilst culture of mESCs in ‘2i’ media results in a reduction of 5mC reflective of their naïve or ground state of pluripotency, the addition of vitC to 2i media causes a further reduction in 5mC, which results in an epigenomic and transcriptomic state more closely resembling the inner cell mass of the blastocyst (5). Human embryonic stem cells behave similarly, with vitC resulting in widespread DNA demethylation in human ESCs (9). VitC also enhances somatic cell reprogramming to more effectively form induced pluripotent stem cells (iPSCs) (10–12). This is in line with the fact that during ESC and induced pluripotent stem cell (iPSC) culture, both TETs and Jumonji histone demethylases are required for effective epigenetic reprogramming (13–17). Indeed, the effects of vitC on DNA demethylation have been demonstrated to occur via TETs (5,6,18,19), and in iPSC reprogramming vitC induces a highly specific reduction in global H3K9me2/3 via KDM3A/B and KDM4B/C, and H3K36me2/3 via KDM2A/B, driving the pre-iPSC to iPSC transition (16,17,20).

There is also evidence that appropriate levels of maternal vitC intake are required for DNA demethylation during germ cell development (21). VitC has been suggested as a promising therapeutic agent in cancer, since it promotes immune cell function, whilst negatively regulating leukemogenesis and tumour invasion (22–24).

Despite a large body of work addressing the impact of vitC on pluripotency, development and cancer, little is known about its effects on transposable elements (TEs). TEs are a diverse set of abundant repetitive sequences that can impact genome regulation in myriad ways (25). A subset of mammalian TEs, including LINE-1 (L1) elements, retains the ability to mobilise within the genome, constituting a mutagenic risk. Notably, both mouse and human pluripotent stem cells support relatively high levels of L1-mediated retrotransposition (26–28). Apart from their mobility, TEs can also act as cis-regulatory elements or trans-acting long non-coding RNAs (29), contribute to 3D genome folding (30), and trigger innate immune responses (31). The question of vitC effect on TEs is particularly relevant given the now common use of vitC in iPSC protocols, and subsequent potential therapies, which may lead to TE activation and subsequent insertional mutagenesis.

Many of the key epigenetic marks that control TE expression, such as H3K9me3 and DNA methylation, can be removed by 2OGXs. We have previously shown that in mESCs, young L1s and endogenous retroviruses (ERVs) of the IAP family are targeted by TET enzymes, but that other epigenetic modifiers counter their activating effect, ensuring these TE families remain repressed (32,33). The inclusion of vitC in the culture media could skew this balance and drive the activation of TEs by enhancing TET activity. Previous work has shown that TEs are activated during the transition of mESCs from serum to 2i+vitC media, but it is unclear what the impact of vitC alone is (34). In another study, IAPs were found to be relatively resistant to vitC-induced demethylation in mESC (5), whilst in human cancer cells, vitC increased the expression of human TEs, including L1, but only in conjunction with DNA methyltransferase inhibition using 5-azacytidine (35).

Here we show that evolutionarily young L1s are activated by vitC treatment in mESCs, through KDM4A/C-mediated H3K9me3 demethylation but independent of TET activity. This did not translate into a measurable increase in L1 retrotransposition, probably due to additional restriction mechanisms. L1 proteins were also upregulated in vitC-treated hESCs, which has potential implications for iPSC derivation and maintenance.

## Results

### VitC upregulates a subset of TEs in 2i-grown mESCs

We first tested the effect of vitC on TE expression in mESCs following a 24h treatment on cells grown in 2i media. 2i media is serum-free, with inhibitors that block the MAPK/ERK and glycogen synthase kinase 3β (GSK3β) pathways, increasing pluripotency gene expression compared to serum-LIF media, for example, and maintaining a ground state of pluripotency (36). We used a selected panel of qRT-PCR primers targeting ERVs, including IAPEz, RLTR4/MuLV, ETn/MusD, MERVL, and evolutionarily young L1s (L1Tf, L1A, L1Gf). The different L1 subfamilies can be distinguished by their variable 5’ UTR sequences, whilst coding regions Orf1 and Orf2 are largely conserved. We observed a robust 2-3 fold upregulation of L1Tf and L1Gf, as well as less substantial changes in expression of some ERV families, including RLTR4/MuLV (Figure 1A). To test whether longer treatments would exacerbate this effect, we performed a time-course experiment for 1 week. Whilst longer treatments resulted in slightly higher levels of TE expression, the most substantial increase occurred within 24h (Figure 1B). We also tested the effect of growth media, and found that adding vitC to mESCs grown in serum/LIF resulted in notably milder effects on TE expression (Figure 1B). We hypothesized that this was due to the reduced 2OG/succinate ratio seen in serum-grown mESCs when compared to 2i-grown cells, as it can limit the activity of 2OGXs (37). However, neither the addition of exogenous 2OG to serum-grown cells or the addition of exogenous succinate to 2i-grown cells substantially altered the action of vitC on L1 expression (Additional file 1: Figure S1A,B).

**Figure 1.**
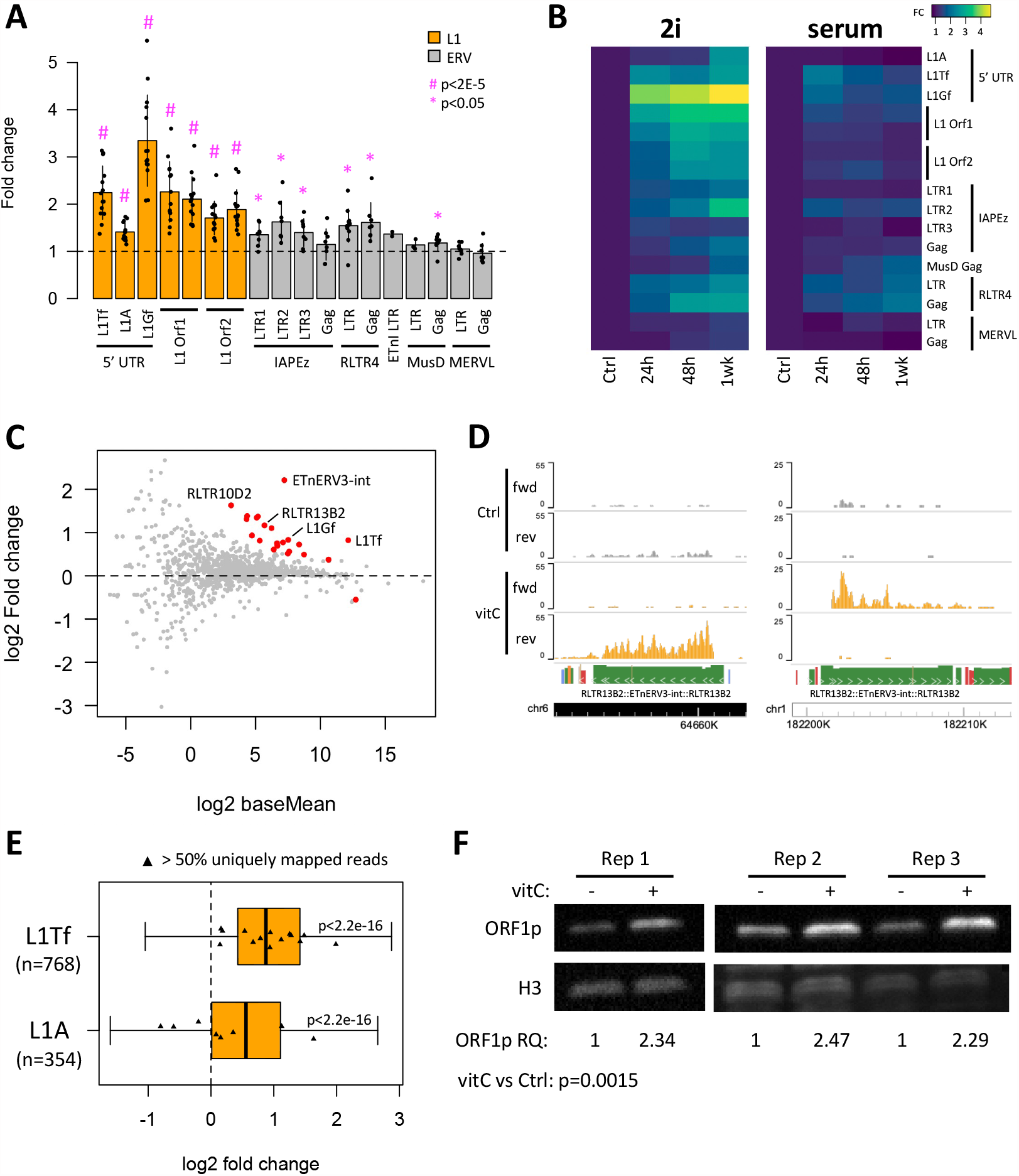
Vitamin C treatment results in upregulation of young L1s and ERVs in mESCs. A) qRT-PCR data showing TE expression fold change in vitC-treated mESC in 2i media compared to untreated cells. P-values are from one-sample t-tests (mu=1) with Benjamini-Hochberg multiple comparisons correction. B) Heatmap from qRT-PCR data showing TE expression fold change in 2i or serum media in vitC-treated versus untreated mESCs at 24 hrs, 48 hrs and after 1 week. C) MA plot for from RNA-seq expression data showing log2 fold change of TE subfamilies following 24hrs vitC treatment of 2i-grown mESCs. Differentially expressed TE subfamilies are coloured red, including L1Tf, L1Gf, RLTR10D2, RLTR13B2 and its internal proviral sequence EtnERV3-int. D) Genome browser snapshots of examples of upregulated RLTR13B2:EtnERV3-int elements in a proviral arrangement. Arrowheads indicate the sense orientation of the elements. E) Boxplot showing log2 fold change for individual L1Tf and L1A elements after mapping of RNA-seq data using SQuIRE. Elements with more than 50% of the reads uniquely mapped are highlighted. P-values are from one-sample t-tests (mu=0) with Benjamini-Hochberg multiple comparisons correction. F) Western blot of ORF1p protein in vitC-containing media compared to controls for three biological replicates. Relative quantification (RQ) values are shown below. P-value is from a t-test.

We decided to focus on the 24h time point (and 2i-grown cells), to minimise potential indirect effects that would confound the analysis, and performed RNA-seq to ask whether expression of other TE families not included in our qPCR panel was modulated by vitC. A total of 150 genes were differentially expressed, most of which were upregulated (Additional file 1: Figure S1C; Additional file 2), as previously observed (5). To analyse TE expression we used SQuIRE, which employs an expectation-maximisation algorithm to assign multi-mapping short reads to individual TE loci (38). A subfamily-level analysis revealed upregulation of several ERVs, most prominently ETnERV3-int (Figure 1C; Additional file 3), a family of the ERV-ß2 group that is distinct from the ETn/MusD group we had initially tested by RT-qPCR (39). Whilst annotated ETnERV3 internal regions are found coupled to either RLTR13B2 or RLTR13A3 LTRs, we only observed upregulation of RLTR13B2, suggesting further subfamily specificity (Figure 1C). Notably, we found that out of ten ETnERV3-int loci with >4-fold upregulation (and mean log2 RPM > -2), eight displayed a proviral arrangement, with RLTR13B2 LTRs flanking both sides of ETnERV3-int fragments (Figure 1D; Additional file 4). Although no ETnERV3 elements are known to be autonomously competent for retrotransposition, four germline mutations due to ETnERV3 insertions have been reported in mouse strains, suggesting recent activity using proteins from other retrotransposons (39).

Our subfamily-level analysis of the RNA-seq data also confirmed that young L1 families (L1Tf, L1Gf) were upregulated upon vitC treatment (Figure 1C; Additional file 3). To ensure that this effect was not confounded by other transcripts harbouring L1 fragments, we looked in more detail at independently transcribed elements larger than 5 kb, which SQuIRE identifies based on the relationship with other transcripts. This analysis suggested that bona fide L1 loci were being activated by vitC (Figure 1E; note that there were insufficient L1Gf loci for a robust analysis). Further subsetting of these elements into those with a high proportion of uniquely mapped reads confirmed that multiple L1Tf loci are upregulated (Figure 1E). L1 activation was also observed at the protein level, as judged by western blot of ORF1p (Figure 1F). These data demonstrate that vitC drives increased TE transcription in 2i-grown mouse ESCs, especially for full-length ETnERV3 elements and putatively active L1s.

### L1 activation by vitC does not require TET activity

We hypothesized that the effects of vitC on TE expression were due to the increased activity of one or more chromatin modifiers of the 2OGX family. In line with this, vitC had no effect on the expression of L1 Orf1 in mESCs treated with DMOG, a pan-inhibitor of 2OGXs, although this result was somewhat confounded by the fact that DMOG alone increased L1 expression (Additional file 1: Figure S2A). We therefore undertook a more specific analysis by targeting candidate 2OGX enzymes.

We have previously shown that both TET1 and TET2, members of the 2OGX family, demethylate young L1s in mESCs (32). To assess whether TET activity at TEs was increased in vitC-treated cells, we measured 5hmC and 5mC levels at selected TEs using targeted oxidative bisulphite sequencing (oxBS-seq) (40). Notably, 5mC levels were reduced at the 5’ UTR of young L1 elements following vitC treatment, concomitant with raised 5hmC levels (Figure 2A). This was also observed at RLTR4/MuLV, but not IAPLTR1a elements (Figure 2A). As the levels of *Tet1* and *Tet2* transcripts did not increase (Additional file 1: Figure S2B), TET activity was likely raised by the co-factor action of vitC. To test whether TETs mediated the effects of vitC on L1 expression, we used *Tet* triple knock-out (TKO) mESCs (i.e., *Tet1*^-/-^;*Tet2*^-/-^;*Tet3*^-/-^) (41), which we confirmed to display no expression of intact *Tet1* or *Tet2* transcripts (Additional file 1: Figure S2C). Surprisingly, treatment of *Tet* TKO mESCs with vitC led to a similar upregulation of young L1s as seen in wild-type (WT) cells (Figure 2B). In contrast, although IAPs also showed no difference in vitC response between the two cell lines (Additional file 1: Figure S2D), the upregulation of other ERVs (e.g., ETnERV3, RLTR4), was visibly impaired in *Tet* TKO cells (Figure 2C). These results show that, whilst TET activity is required for the activation of some ERVs by vitC, it is not necessary for enhancing the expression of L1s, which are the main group of retrotranspositionally active autonomous elements in mESCs.

**Figure 2.**
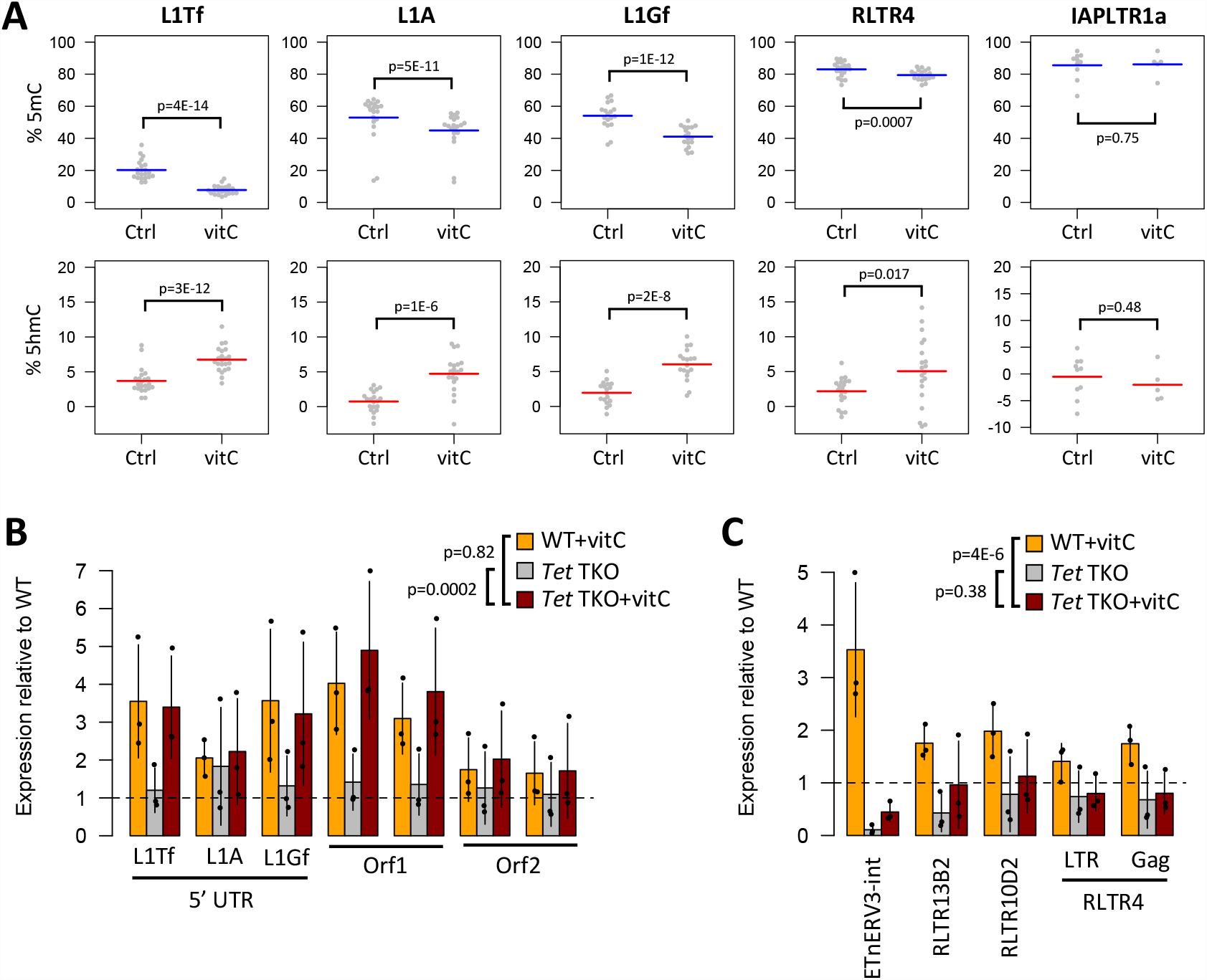
TET activity is dispensable for vitC-mediated L1 upregulation. A) Deep amplicon sequencing of oxBS-treated DNA was used to measure 5mC and 5hmC levels at young L1s in control and vitC-treated mESCs; each data point represents the average value from three biological replicates at a given CpG within the amplicon. P-values are from an ANOVA model. B) qRT-PCR data showing the relative change in expression level (compared to untreated WT cells) for L1s in *Tet* triple knockout (TKO) cells in control and vitC culture conditions. P-values are from an ANOVA model. C) qRT-PCR data as in B, but for ERVs. P-values are from an ANOVA model.

To uncouple the effects of DNA methylation from those of other regulatory marks, we performed experiments in *Dnmt* TKO mESCs (i.e., *Dnmt1*^-/-^;*Dnmt3a*^-/-^;*Dnmt3b*^-/-^) (42), wherein vitC was still able to upregulate L1s and ETnERV3 elements (Additional file 1: Figure S2E). This suggests that other regulatory determinants beyond DNA methylation are key mediators of the vitC response, although at some ERVs DNA methylation may act in concert with other mechanisms. As such, we next investigated whether 2OGX family histone demethylases affected TE expression through vitC-enhanced activity.

### L1 activation by vitC depends on H3K9me2/3 demethylases

TE silencing in mESCs is frequently associated with methylation of H3K9, namely via the TRIM28/KAP1 and HUSH complex pathways (43,44). Notably, global levels of H3K9me2 have been shown to decrease upon vitC treatment in mESCs via the H3K9me1/2 demethylases KDM3A/B (JMJD1A/B), which are 2OGXs (20). We therefore depleted mESCs of both *Kdm3a* and *Kdm3b* using shRNAs, followed by vitC treatment (Additional file 1: Figure S3A). The knockdown did not affect the expression of pluripotency markers, but severely impaired the upregulation of *Dazl* by vitC (Additional file 1: Figure S3A), as previously described (20). In contrast to *Dazl*, L1 elements were still activated by vitC to levels close to those seen in control cells (Figure 3A), implying that other H3K9 demethylases may be involved. Whilst H3K9me2 levels are dramatically reduced genome-wide upon vitC treatment (20), we noted that, compared to the rest of the genome, H3K9me2 levels appear to be better retained at full-length young L1s (Figure 3B). This is in line with KDM3A/B demethylases having a negligible effect on L1 expression after vitC treatment.

**Figure 3.**
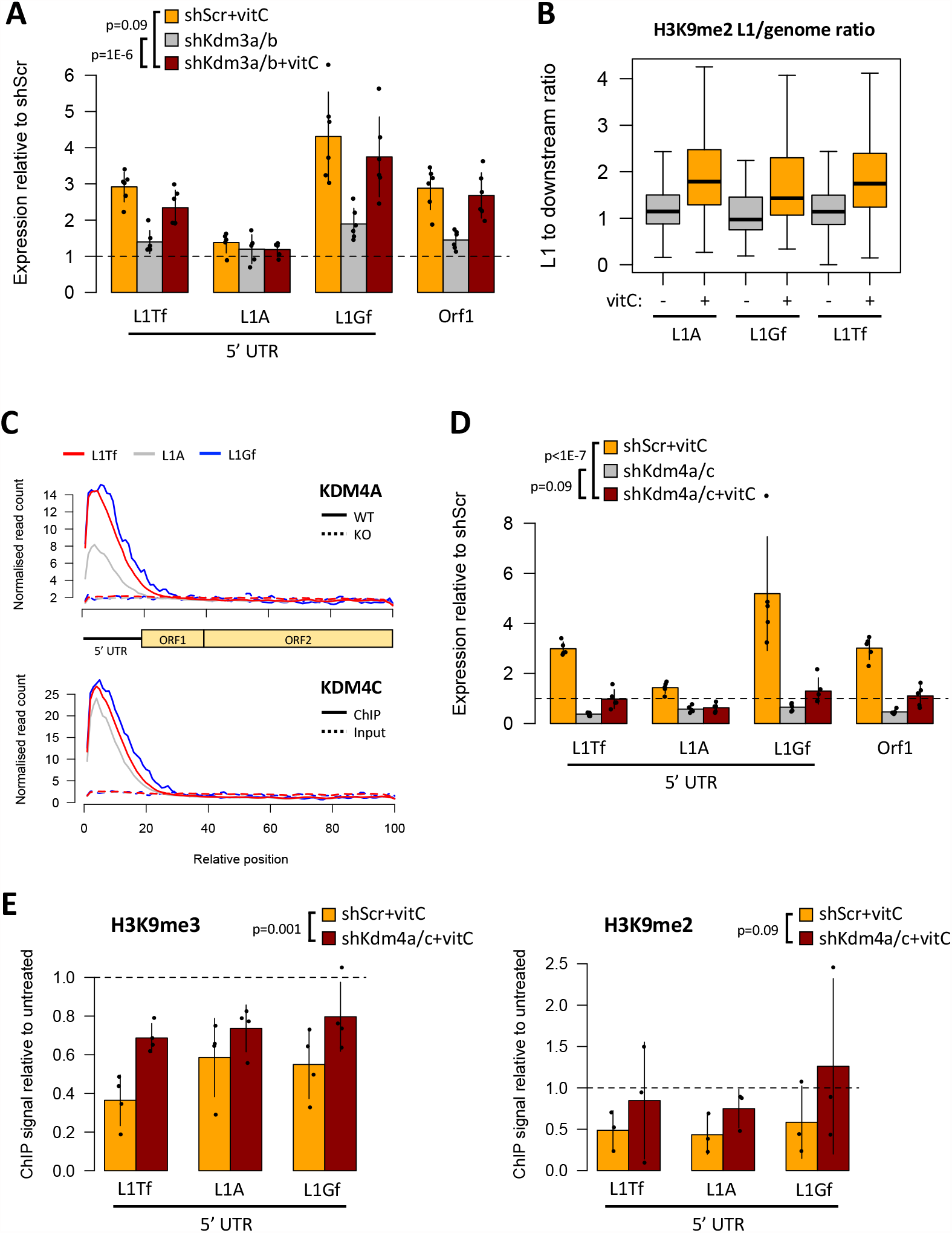
L1 activation by VitC depends on KDM4 H3K9me2/3 demethylases. A) qRT-PCR data showing the expression (relative to untreated shScr) of L1s in sh*Kdm3a/b* double knockdown cells, with and without vitC treatment. B) Using published ChIP-seq data (20), the enrichment of H3K9me2 at young full-length L1s (>5kb; uniquely aligned reads) was compared with that at downstream regions (5kb) from the same elements. In vitC-treated cells (72h), where there is genome-wide H3K9me2 loss (20), L1s preserve more H3K9me2 relative to surrounding regions. C) Cumulative KDM4A (46) and KDM4C (45) ChIP-seq signal at full-length L1s. For KDM4A, the dashed line represents the ChIP signal in *Kdm4a* KO cells; for KDM4C it represents the input signal. D) qRT-PCR data in sh*Kdm4a/c* KD compared to untreated shScr, with and without vitC. E) ChIP-qPCR data for H3K9me3 (left-hand chart) and H3K9me2 (right-hand chart) at the 5’ UTR of young L1s in *Kdm4a/c* KD mESC. The data show ChIP signals in vitC-treated cells normalised to their respective untreated controls. All p-values shown were derived from ANOVA models.

The KDM4/JMJD2 enzymes are key H3K9me2/3 demethylases that play important roles in mESCs and early embryonic development (45,46). The expression of KDM4D is seemingly restricted to the testis (47), but KDM4A, KDM4B and KDM4C are all expressed in mESCs (46). To assess whether KDM4 enzymes could regulate L1 elements, we first reanalysed published KDM4A (46) and KDM4C (45) ChIP-seq data. We included non-unique reads in our alignment and then combined the signal from all copies of the same L1 family. This revealed a strong enrichment of both KDM4A and KDM4C at the 5’ UTR of all three young L1 families, suggesting a putative regulatory role (Figure 3C). L1A elements displayed lower KDM4A enrichment than L1Tf or L1Gf, in correlation with the changes in expression upon vitC treatment. Similar results were obtained with a peak-focused analysis, which also showed that older L1 families were not enriched for either enzyme (Additional file 1: Figure S3B).

Based on these observations, we decided to test for a role of KDM4 demethylases in L1 regulation by vitC. In 2i-grown mESCs, KDM4A and KDM4C display redundant roles in controlling self-renewal, whereas KDM4B appears to be dispensable for this property (46). We therefore depleted mESCs of both *Kdm4a* and *Kdm4c* using shRNAs, which resulted in only minor changes to the expression of pluripotency markers (Additional file 1: Figure S3C). In contrast, *Kdm4a/c* depletion led to a consistent downregulation of L1s, and dramatically limited their activation upon vitC treatment (Figure 3D). Changes in L1 expression were accompanied by corresponding alterations at the level of chromatin, with H3K9me2/3 levels decreasing at the 5’ UTR of L1s upon vitC treatment in control mESCs (Figure 3E). In *Kdm4a/c*-depleted cells the vitC-induced reduction in H3K9me3 levels was less pronounced (Figure 3E), in correlation with the observed L1 expression changes. To confirm these observations, we used an inducible *Kdm4a/b/c* TKO model (46), treating cells with vitC after confirming efficient KO induction (Additional file 1: Figure S3D). Similar to what was seen in shRNA *Kdm4a/c*-depleted cells, *Kdm4a/b/c* TKO cells displayed lower L1 expression than their wild-type counterpart (with the exception of L1A, which was unexpectedly upregulated), and no activation upon vitC treatment, both at the level of RNA (Figure 4A) and protein (Figure 4B). To rule out changes in DNA methylation as a potential confounder, we performed oxBS on *Kdm4a/b/c* TKO cells. Although 5mC levels were higher at L1Tf and L1Gf in cells lacking *Kdm4a/b/c*, in correlation with expression changes, they were reduced (and 5hmC increased) upon vitC treatment with no matching change in expression (Figure 4C). These results show that KDM4A/C activity is necessary for the full activation of L1s upon vitC treatment by controlling H3K9me2/3 deposition.

**Figure 4.**
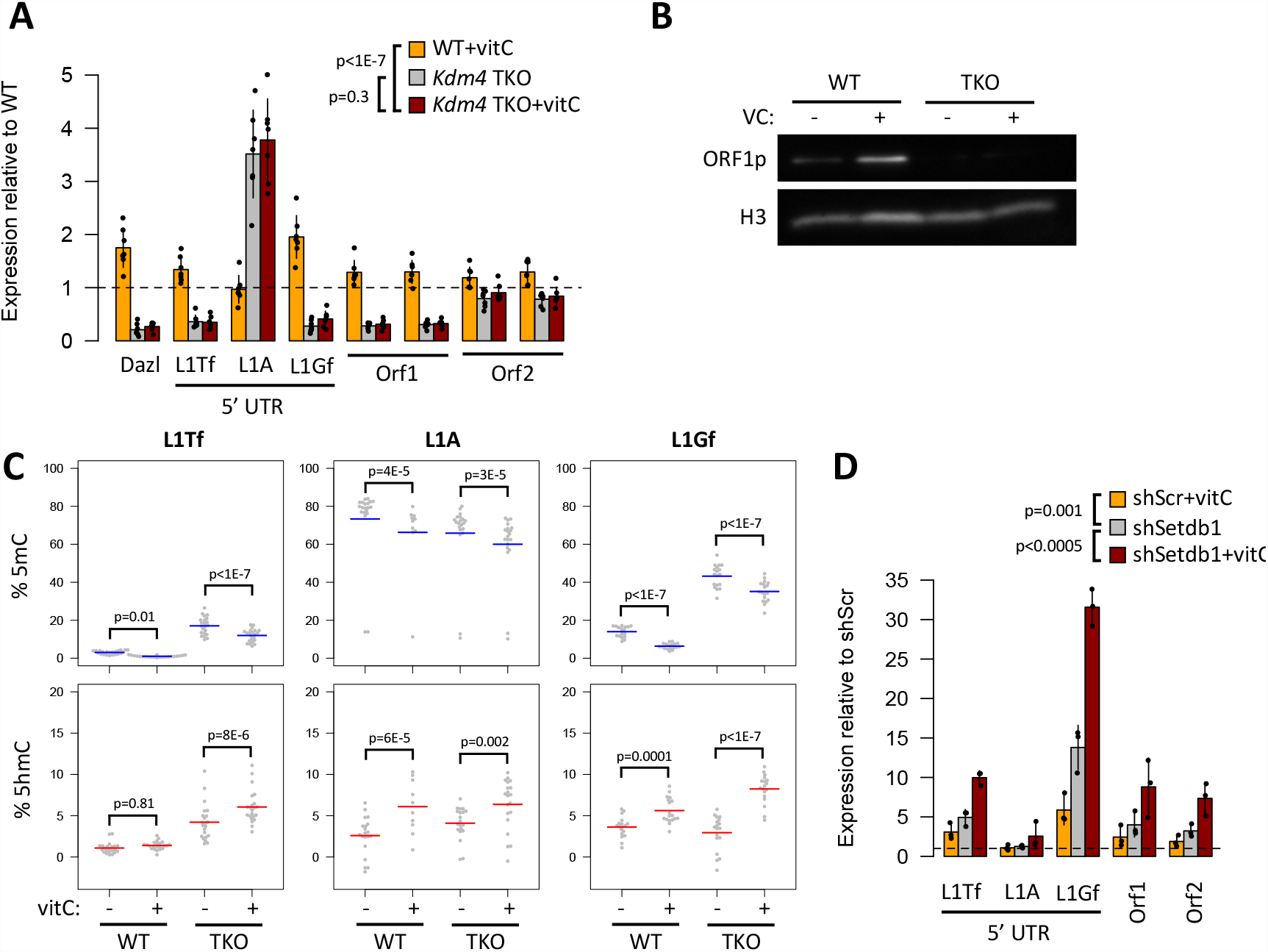
L1 activation by vitC is driven by a reduction in H3K9me2/3. A) qRT-PCR data (relative to untreated WT cells) showing expression of L1s (and *Dazl*) in *Kdm4a/b/c* TKO cells, with and without vitC. B) Western blot showing that L1 Orf1p protein upregulation with vitC is lost in *Kdm4* TKO cells. C) Deep amplicon sequencing from oxBS-treated DNA was used to measure 5mC and 5hmC levels at young L1s in ctrl and vitC-treated *Kdm4* TKO mESCs; each data point represents the average value from three biological replicates at a given CpG within the amplicon. D) qRT-PCR data for L1s in sh*Setdb1* KD mESC, compared to untreated shScr controls, with and without vitC. All p-values shown were derived from ANOVA models.

To confirm that a reduction of H3K9me2/3 methylation was sufficient to increase L1 expression in 2i mESCs, we used shRNA to knockdown *Setdb1*, which encodes for a H3K9 histone methyltransferase that is essential for ERV repression in ESCs and primordial germ cells (48–50). *Setdb1* depletion in mESCs resulted in upregulation of L1 expression (Figure 4D), which is in line with H3K9me2/3 reduction playing a role in vitC-mediated L1 activation. However, when vitC was added to the *Setdb1*-depleted mESCs, L1 expression increased still further (Figure 4D), indicating that additional epigenetic regulators sensitive to vitC remain. Given this high level of L1 expression is not seen in control cells treated with vitC, it is possible that the effects seen in *Setdb1*-depleted mESCs are indirect and/or a result of the long term reduction in H3K9me3 in these cells.

Altogether, these data indicate that loss of H3K9me2/3 is the main driver in vitC-mediated changes in L1 expression.

### No evidence of increased L1 retrotransposition upon vitC treatment

The upregulation of transcripts and protein from full-length, evolutionarily young L1s, raised the possibility that vitC treatment leads to increased somatic L1-mediated retrotransposition in mESCs. To test this, we aimed to detect endogenous de novo TE insertions using a target-sequencing approach in combination with custom-made sequencing adapters (Additional file 5) – similar to SIMPLE (51) and ME-Scan (52). In a first experiment, we analysed four independent mESC clones treated with vitC for 13 days, and compared them with untreated controls (Figure 5A). After data processing with TEBreak, only two candidate de novo TE insertions were identified, one from a control sample (L1) and the other from a vitC-treated sample (SINE B2). An additional 58 insertions (57 L1, 1 SINE B1) were exclusive to just one of the four clones, but present in both control and vitC-treated samples, suggesting they were already present at the start of the experiment.

**Figure 5.**
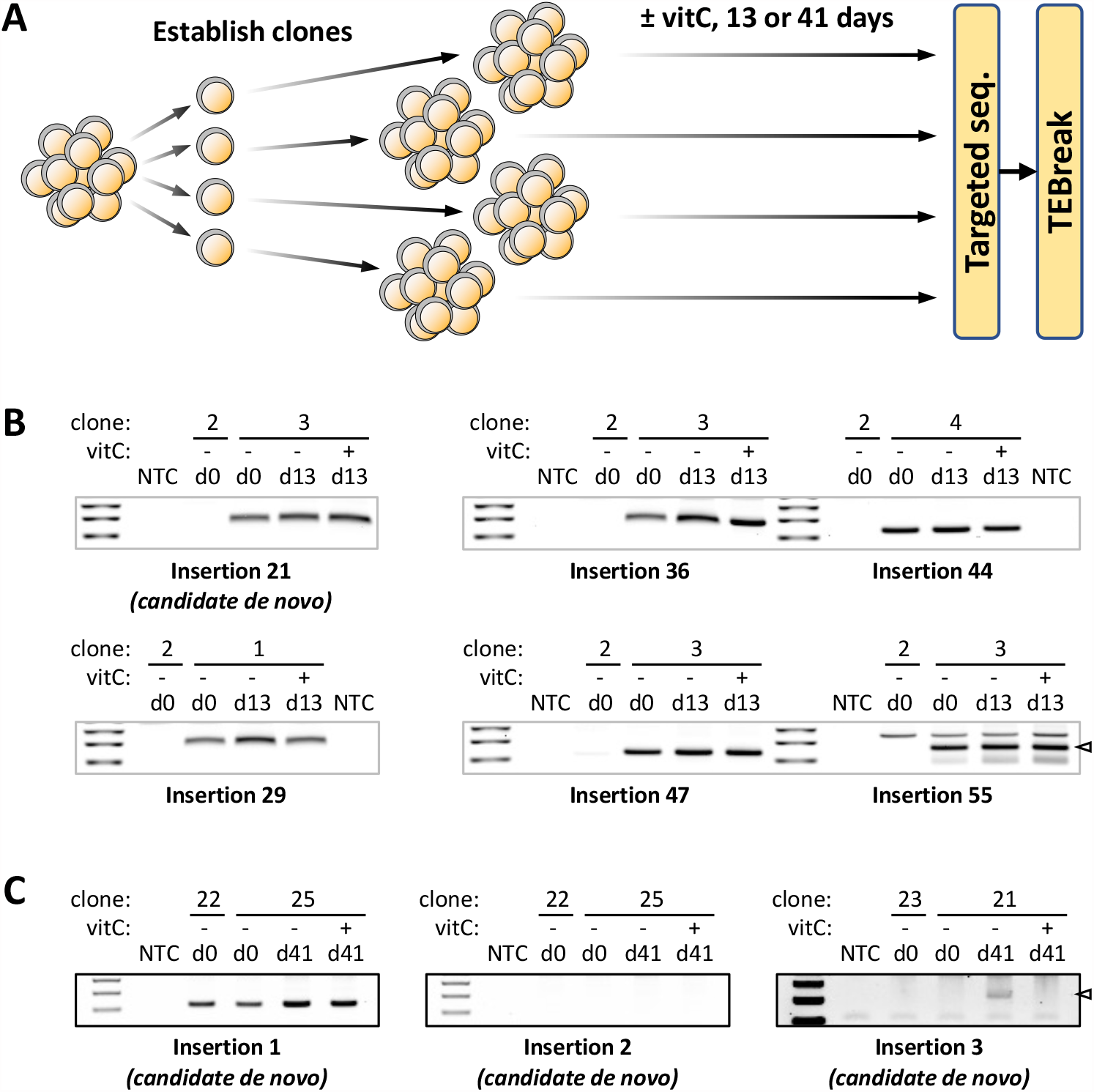
No evidence of increased L1 retrotransposition upon vitC treatment. A) Schematic showing the design of the retrotransposition experiment undertaken. Four clonal cell populations were grown in 2i media or 2i+vitC for 13 or 41 days, followed by L1 targeted sequencing analysis and processing by TEBreak to identify de novo retrotransposition events. B) Agarose gel electrophoresis detection of candidate TE insertions following PCR on samples from the 13-day experiment. All 6 analysed insertions were detected in both control and vitC samples of the expected clone, including a candidate de novo insertion. C) As in B, but for the 41-day experiment. One bona fide de novo insertion was identified that could be validated by PCR, which occurred in the non-vitC control condition. NTC: no template control.

To scrutinise the reliability of these results we undertook PCR-based validations. We analysed the potential de novo L1 insertion and 5 other insertions with a highly skewed read count towards either the control or the vitC-sample that could be due to an artefactual sequencing call. However, all of the analysed insertions (n=6) were detected in both control and vitC-samples of the expected clone, including the candidate de novo insertion, which suggests it had been detected in only one sample due to sequencing drop-out (Figure 5B).

We then performed a second experiment with vitC treatment for 41 days, to provide additional time for de novo insertions to occur and become established within the population by genetic drift. Similar to the first experiment, however, only three insertions were putatively de novo, two in control samples and one in a vitC-treated sample. Only one of these insertions (in a control sample) was validated by PCR (Figure 5C). Finally, we maximised our sensitivity by reducing the stringency of our analysis to include insertions with support from 1-4 reads, although this comes at the cost of decreased specificity. We identified 350 putative de novo L1 insertions in control samples and 301 in vitC-treated samples, again finding no evidence that vitC treatment leads to increased L1-mediated retrotransposition in mouse ESCs.

It remains possible that vitC treatment increases the number of low frequency L1 insertions within the cell population, which would be under our detection limit.

Nonetheless, our results demonstrate that this does not translate into a significant shift in the L1 makeup of the ESC population as a whole.

### VitC activates human L1s in hESCs

In human pluripotent cell culture, vitC increases the efficiency of reprogramming in iPSC (11). The mechanism behind this is unclear, although in primed hESCs vitC leads to widespread DNA demethylation and is associated with decreased H3K9 methylation (9). Since histone demethylases appear to be key in mediating the effects of vitC in mESCs, we first explored whether they may also be involved in human ESCs. Using ENCODE data, we found that, similar to mESC, KDM4A is bound at the 5’ UTR of young L1s in hESCs (Figure 6A). We therefore treated primed hESCs with vitC but, differently to the mouse, observed no transcriptional effect on L1 expression at 24h or 48h (Figure 6B). However, we did observe an increase in ORF1p levels in both primed and naïve hESCs (Figure 6C), again seemingly independent of transcriptional changes (Additional file 1: Figure S4A). Since measures of L1 expression by RT-qPCR can be confounded by elements overlapping genes and pervasive transcription, we performed RNA-seq on vitC-treated primed hESCs to better dissect L1 expression. We found no overall activation of young L1 subfamilies after analysis with SQuIRE (Figure 6D). We obtained the same result with TEtranscripts (Additional file 1: Figure S4B) and TExP (Additional file 1: Figure S4C), which is a tool to remove the confounding influence of pervasive transcription (53). We then extended the SQuIRE analysis to individual L1Hs, L1PA2 and L1PA3 full-length elements, but found no evidence of substantial L1 upregulation after vitC treatment (Figure 6E).

**Figure 6.**
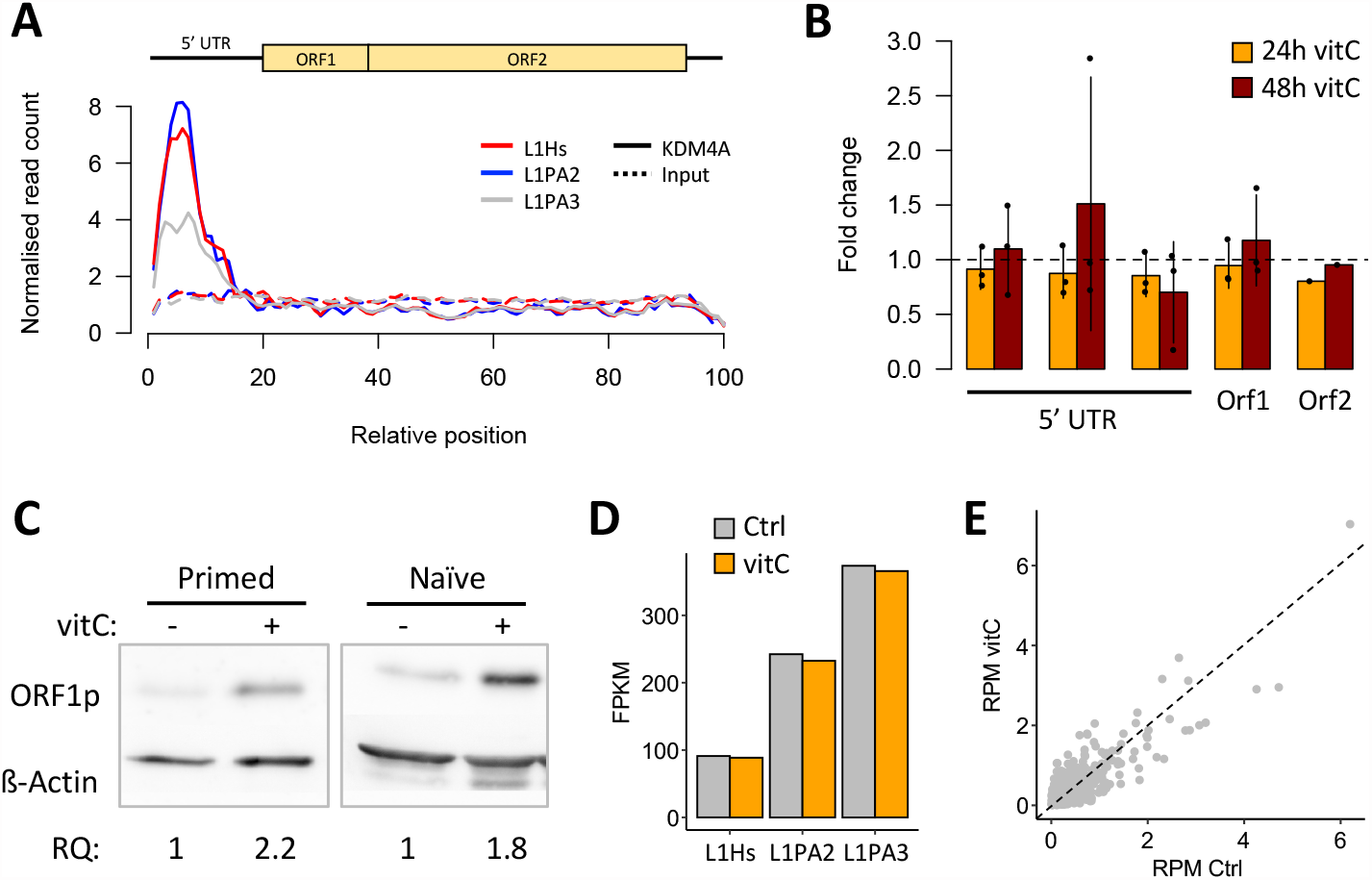
VitC activates human L1s in hESCs. A) Alignment of ChIP-seq data for KDM4A in human ESCs (ENCODE data) showing that KDM4A is highly enriched at the 5’ UTR of young L1s (L1Hs, L1PA2). B) qRT-PCR data showing L1 expression in primed hESCs (H9 NK2) following vitC treatment for 24 or 48 hrs, compared to untreated controls. Multiple 5’ UTR primers were used. C) Western blot for human ORF1p in primed (H9, left panel) and naïve (H9 NK2, right panel) hESCs. Relative quantification (RQ) values are shown below. D) Analysis of RNA-seq data in control and 24h vitC-treated primed hESCs by SQuIRE, showing subfamily level expression of young L1s. E) Expression of individual full-length L1 loci (from L1Hs, LPA2 or L1PA3 families) in control and vitC-treated cells, analysed by SQuIRE. Note that the values (reads per million) are in a linear scale, not log-transformed.

These data demonstrate that vitC upregulates L1 proteins in hESC, and thus potentially retrotransposition therein. Whilst we cannot exclude the possibility that a specific upregulated L1 transcript contributes to most protein signal, our data suggest that vitC increases ORF1p expression in hESCs via a distinct mechanism to that in mESC, acting at the post-transcriptional level.

## Discussion

During cellular reprogramming, the epigenome undergoes significant waves of DNA and histone demethylation, promoting the formation of a pluripotent epigenetic profile and naïve state (54). Over the past few decades, tissue culture regimens have been revolutionised, allowing this process to be reconstructed in vitro, resulting in the development of ground-state ESC culture, and, perhaps more importantly, the induction of pluripotency from somatic cells (55–57). VitC has been well characterised as a required co-factor for the function of two key enzyme classes for this process, TETs and Jumonji histone demethylases, and vitC addition to culture media increases the efficiency of epigenetic reprogramming (5–8). However, the downstream effects of vitC-mediated epigenetic reprogramming have not been fully characterised.

Here, we found that young L1 families and several ERVs are derepressed following vitC treatment, resulting in their increased transcription in mESCs and increased L1 Orf1p in mouse and human ESCs. We show that DNA methylation reprogramming is insufficient for this process at L1 loci, but that TETs are required for ERV transcriptional upregulation. At L1s, vitC-mediated enhancement of KDM4A/C histone demethylase activity is largely responsible for L1 overexpression by reducing H3K9me2/3 levels.

L1 activity was specific to younger L1 classes in mouse, namely L1Gf (1.45 MYR old), L1A (1.33 MYR old), and L1Tf (0.58 MYR old) (58–60). The expansion of these elements in the mouse genome was coincident with a reduction in TRIM28/KAP1 binding, which mainly targets older L1s (58). Nonetheless, in conjunction with the HUSH complex, TRIM28 still plays a key role in silencing young L1s (43), which is in agreement with our observations that H3K9me2/3 modulation by vitC impacts L1 expression. The L1A class behaved slightly differently to L1Gf and L1Tf in response to VitC, exhibiting a generally a more dampened response. This could be due to differences in the regulation of L1A copies, since H3K9me3 deposition at these loci is enacted by SUV39H1/2 (61) instead of SETDB1, which may result in differences in the steady-state levels of H3K9me3 across L1 families.

The possibility that vitC treatment could lead to increased somatic L1 retrotransposition in ESCs has far reaching concerns, since a large number of tissue culture protocols include vitC in their media, meaning that many lines used in research may be subject to increased levels of L1-mediated insertional mutagenesis. Our data suggest that, as far as we could detect, vitC does not substantially increase the likelihood that L1 elements will move. Whilst it is possible that novel insertions occurred that we did not detect, their inability to overtake the culture indicates that they are not of practicable concern, and are certainly not drastically affected by vitC. Increased L1 expression could still have other destabilising effects on genomic integrity, such genetic alterations mediated by the Orf2p-encoded endonuclease. Additionally, L1 proteins could mediate retrotransposition of other elements not included in our assay, such as ETnERV3, which were highly upregulated in response to vitC and display evidence of recent mobility in the mouse genome (39). L1 transcripts have also been shown to act as long non-coding RNAs in mESCs, silencing 2 cell-stage genes and increasing the expression of ribosomal RNA (62), but higher L1 transcription after vitC treatment is unlikely to have further effects on either of these processes. It remains to be tested whether higher Orf1p expression in hESCs results in higher retrotransposition rates therein, as well as what the specific vitC-dependent mechanisms are that lead to its post-transcriptional upregulation.

Our work demonstrates that expression of young L1 elements and some ERVs are directly activated by vitC through epigenetic mechanisms in ESCs. Given the myriad roles and impact of TEs on genome regulation and integrity, there may be downstream consequences to be uncovered. Additional relationships between environmental cues and TE activity may also exist in this and other cell types.

## Methods

### Mouse ESC culture

E14 mESCs were used for all experiments unless stated otherwise. *Tet1/2/3* triple KO cells are as described in (41) and *Dnmt1/3a/3b* triple KO cells in (42).

*Jmjd2a/b/c*^f/f^;Rosa26::CreERT2 conditional KO cells (46) were supplemented with 500nM 4-hydroxytamoxifen for 10 days to induce Cre recombinase expression. ESCs were grown in feeder-free conditions, on gelatinized cell culture dishes with either ‘serum media’, containing DMEM-F12 with 15% FBS and 1000 U/mL ESGRO LIF (Millipore), or serum-free ‘2i’ medium (57) comprising 50% DMEM/F-12 1:1 (Invitrogen), 50% neurobasal media (Invitrogen) supplemented with N2 (Invitrogen), B27 (Invitrogen), β-mercaptoethanol (Gibco), 0.1 mM non-essential amino acids (Gibco), 2 mM GlutaMAX (Gibco), 1 mM sodium pyruvate (Gibco), 1000 U/mL ESGRO LIF (Millipore), 1 μM PD-035901 (Biovision) and 3 μM CHIR-99021 (Biovision). For vitC supplementation, vitamin C (Sigma A8960-5G) was dissolved in molecular grade water and added to a final concentration of 100 µg/ml. Treatment of cells with dimethyl alpha-ketoglutarate (4 mM final concentration; Sigma), dimethyl succinate (4 mM final concentration; Sigma) or DMOG (1 mM; Sigma) was carried out for 24 hours. For shRNA-mediated knockdowns, mESCs were infected with lentiviral particles carrying pLKO.1 constructs harbouring gene-specific shRNAs or a non-targeting sequence (Additional file 6). After 24 hours, cells were selected with 10 μg/ml puromycin or 50 μg/ml hygromycin for 3–9 days.

### Human ESC culture

Primed H9-NK2 cells containing doxycycline-inducible KLF2 and NANOG transgenes coupled to Venus were maintained in conventional medium (KSR/FGF) comprised of DMEM/F-12 (Sigma Aldrich) with 20% KSR (ThermoFisher Scientific) and 10 ng per ml basic fibroblast growth factor (bFGF; Peprotech), supplemented with 2mM l-glutamine (ThermoFisher Scientific), 100 μM 2-mercaptoethanol (2ME) (ThermoFisher Scientific), 1% MEM non-essential amino acids (ThermoFisher Scientific), and 50 mg per ml Penicillin-Streptomycin. Cultures were passaged every 5-6 days as small clumps by dissociation with a buffer containing 1 mg per ml Collagenase IV (ThermoFisher Scientific), 0.025% Trypsin (ThermoFisher Scientific), 1 mM CaCl2 and KSR at a final concentration of 20% in PBS. Medium was changed daily. Primed H9 cells were maintained in HFF-conditioned media (HFF-CM) supplemented with 8 ng/ml of β-FGF (MILTENYI BIOTE) on Matrigel (BD, Biosciences) coated surfaces cell lines in a humidified 5% CO2 incubator at 37ºC. Cells were passaged when cultures reach 70– 100% confluence using Collagenase IV (GIBCO) in presence of Rho-associated kinase inhibitor (ROCKi) (Y-27632, Calbiochem). Naïve H9 NK2 cells were grown in feeder-free conditions and 5% O2, plated on diluted growth factor reduced Matrigel (1:100 diluted in DMEM/F12, Invitrogen) with t2iL+Gö (DMEM/F12 and neurobasal media 50:50, Invitrogen), supplemented with N2 (Invitrogen), and B27 (Invitrogen), 1% L-glutamine, 1% β-mercaptoethanol, 0.5x penicillin/streptomycin, 1 µM PD0325901 (Stem Macs, Miltenyi Biotec), 1 µM CHIR99021 (Stem Macs, Miltenyi, Biotec), 2 µM Gö6983 (Sigma-Aldrich), 20 ng/ml LIF (Millipore). Cells were passaged every 3-4 days using Accutase in presence of ROCKi (Y-27632, Calbiochem). VitC supplementation was performed as for mESCs.

### RT-qPCR and RNA-seq

RNA was extracted with Quick-DNA/RNA Miniprep Kit (Zymo Research) and DNAse treated with the TURBO DNA-free Kit (Ambion). cDNA was then prepared with SuperScript III Reverse Transcriptase (Invitrogen). Target sequences (primer list in Additional file 6) were amplified with MESA BLUE qPCR MasterMix Plus (Eurogentec) or KAPA SYBR FAST Roche LightCycler 480 qPCR Master Mix (Kapa Biosystems, KK4611) and quantified on a Light-Cycler 480 (Roche). Prior to RNA-seq library construction, 10-100 ng of total RNA was treated with the NEBNext rRNA Depletion Kit. Library construction was performed with the NEBNext Ultra II Directional RNA Library Prep Kit for Illumina, according to the manufacturer’s protocol.

### Western blotting

Mouse ESCs were washed with ice-cold PBS and lysed in Laemmli buffer (300 mM Tris/HCl pH 6.8, 10 % SDS, 50 % Glycerine, 0.05% Bromophenol Blue and 5% B-Mercaptoethanol), followed by sonication. Human ESCs were lysed with 50-100 µL of RIPA buffer (Sigma) supplemented with 1x Complete EDTA-free Protease Inhibitor cocktail (Roche), PMSF (Sigma), 0.25% β-mercaptoethanol (Sigma), followed by centrifugation. Samples were denatured at 95 ºC for 5 minutes before separation by SDS-PAGE and transfer into nitrocellulose membranes. Membranes were blocked with 5% milk and 1% BSA in PBS-T or with 5% milk in TBS-T for 30-60 min at room temperature and incubated overnight with Rb-anti-mORF1p (1:5,000, kind gift from Alex Bortvin), Rb HRP-conjugated anti-Histone H3 antibody (ab21054), Rb anti-hORF1p (1:1000, kind gift from Oliver Weichenrieder), or anti-actin (1:10,000, Sigma, A2228).

After washing with PBS-T or TBS-T, membranes were incubated with peroxidase-conjugated anti-rabbit (1:5,000 Sigma-Aldrich, or 1:1,000 Cell Signalling) or anti-mouse (1:1,000 Cell Signalling) IgG for 1 h, washed with PBS-T or TBS-T and exposed to ECL reagent (Sigma-Aldrich, WBKLS0500) or Clarity ECL Western Blotting Substrate (BioRad). Signals were visualized using X-ray film, a ChemiDoc MP Imaging System (Bio-Rad), or an ImageQuant LAS4000.

### Chromatin Immunoprecipitation

Cells were fixed with 1% formaldehyde for 12 minutes in PBS, then quenched with glycine (final concentration 0.125 M), washed twice with ice-cold PBS and flash frozen. Cells were first lysed in lysis buffer 1 (5 mM HEPES, 140 mM NaCl, 2.5% glycerol, 0.5% IGEPAL, 0.25% Triton-X, 1 mM EDTA) with complete protease inhibitors (PI, Roche) at 4ºC for 30 min, then in lysis buffer 2 (10 mM Tris-HCl pH8, 200 mM NaCl, 1 mM EDTA, PI) at 4ºC for 30 mins, and resuspended in sonication buffer (10 mM HEPES, 0.5% SDS, 1 mM EDTA, PI). Chromatin was sonicated using a Bioruptor Pico (Diagenode) to an average size of 200-700 bp, then pre-cleared with blocked (with 3 mg/ml salmon sperm DNA and 7 mg/ml BSA for 1h) Protein G magnetic Dynabeads for 2h at 4ºC. After recovering the supernatant, 1 µg of chromatin was removed as input.

Immunoprecipitation was performed overnight using 10 µg of chromatin diluted in ChIP dilution buffer (0.01% SDS, 1.1 % Triton X-100, 1.2 mM EDTA, 16.7 mM Tris-HCl, 167 mM NaCl, PI, pH 8.1) and 2.5 µg of mouse antibodies to H3K9me2 (Diagenode C15410060) and H3K9me3 (Diagenode C15410193). Samples were incubated with blocked Protein G Dynabeads for 4 h, then washed twice with wash buffer A (50 mM Tris-HCl pH 8, 150 mM NaCl, 1 mM EDTA, 0.1% SDS, 1% NP-40, 0.5% deoxycholate, PI), once with wash buffer B (as wash buffer A but with 500 mM NaCl), and once with wash buffer C (50 mM Tris-HCl pH 8, 250mM LiCl, 1 mM EDTA, 1% NP-40, 0.5% deoxycholate, PI). DNA was eluted elution in 0.1 M NaHCO_3_ + 1% SDS, for 40 mins at 65ºC, treated with RNAse A (10 mg/ml) for 15 mins at 37ºC, and with proteinase K (20 mg/ml) overnight at 65ºC. Final DNA purification was performed using the GeneJET PCR Purification Kit (Thermo Scientific, K0701) and eluted in 80 µl of elution buffer. DNA was diluted 1/10 and analysed by qPCR using KAPA SYBR FAST Master Mix (primer list in Additional file 6).

### Bisulphite/oxidative bisulphite sequencing

Precipitated DNA (without glycogen) was resuspended in water and further purified using Micro Bio-Spin columns (Bio-Rad), after which half of the DNA was oxidised with 15 mM KRuO4 (Alpha Aesar) in 0.5 M NaOH for 1 h. The EpiTect Bisulfite kit (QIAGEN) was used to carry out bisulphite conversion of both DNA fractions, followed by a two-step PCR amplification: the first PCR amplifies the region of interest (primers in Additional file 6) and adds part of the sequencing adaptors; the second PCR (on pooled amplicons) completes the adaptors and adds sample barcodes, allowing for multiplexing (40). Paired-end sequencing of pooled samples was carried out on an Illumina MiSeq.

### Somatic L1 insertion detection

The general strategy is described in Additional file 5. Approximately 0.5-1 µg of genomic DNA was sheared using a Covaris E220 Focused Sonicator, in microtubes AFA Fiber Snap-Cap 130, in 120µl of final volume and using the following parameters: Peak Incident Power (W) 50; Duty Factor 20%; Cycles per Burst 200; and Shearing Time 65 sec. The sheared DNA was cleaned up and concentrated using 132 µl AMPure XP Beads (Beckman Coulter), then resuspended in 20 µl final of Illumina TruSeq DNA LT Nano library prep kit resuspension buffer. End-repair was performed using the kit’s protocol and, after beads-clean up, the samples were eluted in 15 µl of resuspension buffer. Size selection was performed by electrophoresis in 2% agarose gel. Two gel excisions were recovered aiming for fragments 450-350bp and 350-250 bp, and the DNA purified by phenol:chlorophorm extraction and ethanol precipitation, resuspending each fragment in 8.75 µl of resuspension buffer. A-tailing, adapter ligation and PCR enrichment were performed using the Illumina TruSeq DNA LT Nano library prep kit protocol and reagents, with modifications. The Illumina adapters were replaced by the homemade ones (see Additional file 5) and the PCR was set for 12 cycles. The shorter fragment were used for enriching 3’ junctions, by using a primer cocktail including the primers TS-Fw and Adapter_mL1_3end (Additional file 6) at 50 µM each. The larger fragment were used for enriching 5’ junctions, by using a primer cocktail including the primer TS-Fw at 50 µM and several primers targeted to the 5’ ends of L1s (Additional file 6) at 8.33 µM. Libraries were pooled and sequenced in an Illumina MiSeq Instrument using a V3 600cycles Reagent Kit, including an ∼20% PhiX spike-in library, and using custom primers (see Additional file 5).

### PCR validation of somatic insertions

PCR validations were performed using MyTaq DNA polymerase (Bioline), including 20 pmol of each primer (Additional file 6), 1 U of enzyme and 10 ng of genomic DNA input, in a final reaction volume of 20 µL and with the following cycling conditions: (95ºC, 2min)x1; (95ºC, 15sec; 60ºC, 20sec; 72ºC, 30sec)x30; (72ºC, 5min; 4ºC, hold)x1.

### High-throughput sequencing data processing

RNA-seq data was analysed using SQuIRE (38). Mapping was done to the mm10 or hg38 genome assemblies using SQuIRE’s Map tool, and read counts per gene, TE or TE subfamily obtained using SQuIRE’s Count tool. These counts were then used for differential expression analysis using DESeq2 (63). Only mouse L1s larger than 5kb and classified as ‘individual TE loci’ under SQuIRE’s Flag script were considered. Wiggle plots were generated using SQuIRE’s Draw tool. Human L1 expression was also quantified using TEtranscripts (64) and TeXP (53).

Reads from published ChIP-seq data (see data and code availability statement) were trimmed using Trim_Galore! and mapped to the mm10 or hg38 genome assemblies using bowtie2 (65) with default settings, which assigns non-unique reads to one of the mapping hits at random. These ambiguously mapped reads were kept for generating average profile plots over full-length (>5kb) L1 elements and for the detection of ‘ambiguous’ KDM4A peaks. Unique reads (MAPQ≥2) were used for H3K9me2 data analysis, and the detection of ‘unique’ KDM4A peaks. Peak detection was performed using MACS2 (66).

OxBS data were aligned with Bismark (67) to a custom genome containing the respective PCR amplicon sequences; only CpGs covered by at least 100 reads were used to calculate 5mC/5hmC levels.

Somatic TE insertions were mapped using TEBreak (https://github.com/adamewing/tebreak) with options --max_disc_fetch 0, --min_maxclip 20, -c 20000, -p 120, --min_disc_reads 0, and --min_split_reads 1 or 4.

## Declarations

### Availability of data and materials

RNA-seq data have been deposited in NCBI’s Gene Expression Omnibus under accession number GSE238224. Publicly available ChIP-seq data were used for H3K9me2 (GEO:GSE84009), KDM4C (GEO:GSE93721), and mouse (GEO:GSE64252) and human (ENCODE:ENCSR000AVC) KDM4A. All remaining data and code associated with the manuscript is available via the GitHub repository https://github.com/MBrancoLab/Cheng_2023_vitC.

## Authors’ contributions

K.C.L.C. and J.M.F performed mESC cell culture, shRNA-mediated knockdowns, RT-qPCR, ChIP, western blots, BS/oxBS assays and RNA-seq. F.J.S.-L. performed assay for detection of somatic L1 insertions. M.G.-C. and H.P. performed experiments on human ESCs, with support from J.L.G.-P. and G.F.. D.T. contributed to ChIP experiments.

W.R.Y. analysed mESC RNA-seq data with SQuIRE, supported by K.H.B.. B.I. and S.S. contributed to experiments on the effects of metabolites and *Kdm3a/b* knockdown. K.A. and K.H. generated the *Kdm4* TKO mESCs. A.E. analysed data from L1 insertion assay.

M.R.B. designed and coordinated the project, performed bioinformatic analyses, and contributed to RNA-seq and somatic L1 insertion experiments. J.M.F and M.R.B wrote the manuscript with all other authors.

## Acknowledgements

We thank Guoliang Xu for the *Tet* TKO mESC line, Alex Bortvin for the anti-mORF1p antibody, and Oliver Weichenrieder for the anti-hORF1p antibody.

## References

1. Gut P, Verdin E. The nexus of chromatin regulation and intermediary metabolism. Nature. 2013 Oct 24;502(7472):489–98.

2. Dai Z, Ramesh V, Locasale JW. The evolving metabolic landscape of chromatin biology and epigenetics. Nat Rev Genet. 2020 Dec;21(12):737–53.

3. Carey BW, Finley LWS, Cross JR, Allis CD, Thompson CB. Intracellular α-ketoglutarate maintains the pluripotency of embryonic stem cells. Nature. 2015 Feb 19;518(7539):413–6.

4. Thienpont B, Steinbacher J, Zhao H, D’Anna F, Kuchnio A, Ploumakis A, et al. Tumour hypoxia causes DNA hypermethylation by reducing TET activity. Nature. 2016 Sep 1;537(7618):63–8.

5. Blaschke K, Ebata KT, Karimi MM, Zepeda-Martínez JA, Goyal P, Mahapatra S, et al. Vitamin C induces Tet-dependent DNA demethylation and a blastocyst-like state in ES cells. Nature. 2013 Aug 8;500(7461):222–6.

6. Young JI, Züchner S, Wang G. Regulation of the epigenome by vitamin C. Annu Rev Nutr. 2015 May 6;35:545–64.

7. Minor EA, Court BL, Young JI, Wang G. Ascorbate induces ten-eleven translocation (Tet) methylcytosine dioxygenase-mediated generation of 5-hydroxymethylcytosine. J Biol Chem. 2013 May 10;288(19):13669–74.

8. Monfort A, Wutz A. Breathing-in epigenetic change with vitamin C. EMBO Rep. 2013 Apr;14(4):337–46.

9. Chung T-L, Brena RM, Kolle G, Grimmond SM, Berman BP, Laird PW, et al. Vitamin C promotes widespread yet specific DNA demethylation of the epigenome in human embryonic stem cells. Stem Cells. 2010 Oct;28(10):1848–55.

10. Stadtfeld M, Apostolou E, Ferrari F, Choi J, Walsh RM, Chen T, et al. Ascorbic acid prevents loss of Dlk1-Dio3 imprinting and facilitates generation of all-iPS cell mice from terminally differentiated B cells. Nat Genet. 2012 Mar 4;44(4):398–405, S1.

11. Esteban MA, Wang T, Qin B, Yang J, Qin D, Cai J, et al. Vitamin C enhances the generation of mouse and human induced pluripotent stem cells. Cell Stem Cell. 2010 Jan 8;6(1):71–9.

12. Chen J, Guo L, Zhang L, Wu H, Yang J, Liu H, et al. Vitamin C modulates TET1 function during somatic cell reprogramming. Nat Genet. 2013 Dec;45(12):1504–9.

13. Doege CA, Inoue K, Yamashita T, Rhee DB, Travis S, Fujita R, et al. Early-stage epigenetic modification during somatic cell reprogramming by Parp1 and Tet2. Nature. 2012 Aug 30;488(7413):652–5.

14. Costa Y, Ding J, Theunissen TW, Faiola F, Hore TA, Shliaha PV, et al. NANOG-dependent function of TET1 and TET2 in establishment of pluripotency. Nature. 2013 Mar 21;495(7441):370–4.

15. Gao Y, Chen J, Li K, Wu T, Huang B, Liu W, et al. Replacement of Oct4 by Tet1 during iPSC induction reveals an important role of DNA methylation and hydroxymethylation in reprogramming. Cell Stem Cell. 2013 Apr 4;12(4):453–69.

16. Chen J, Liu H, Liu J, Qi J, Wei B, Yang J, et al. H3K9 methylation is a barrier during somatic cell reprogramming into iPSCs. Nat Genet. 2013 Jan;45(1):34–42.

17. Wang T, Chen K, Zeng X, Yang J, Wu Y, Shi X, et al. The histone demethylases Jhdm1a/1b enhance somatic cell reprogramming in a vitamin-C-dependent manner. Cell Stem Cell. 2011 Dec 2;9(6):575–87.

18. Dickson KM, Gustafson CB, Young JI, Züchner S, Wang G. Ascorbate-induced generation of 5-hydroxymethylcytosine is unaffected by varying levels of iron and 2-oxoglutarate. Biochem Biophys Res Commun. 2013 Oct 4;439(4):522–7.

19. Yin R, Mao S-Q, Zhao B, Chong Z, Yang Y, Zhao C, et al. Ascorbic acid enhances Tet-mediated 5-methylcytosine oxidation and promotes DNA demethylation in mammals. J Am Chem Soc. 2013 Jul 17;135(28):10396–403.

20. Ebata KT, Mesh K, Liu S, Bilenky M, Fekete A, Acker MG, et al. Vitamin C induces specific demethylation of H3K9me2 in mouse embryonic stem cells via Kdm3a/b. Epigenetics Chromatin. 2017 Jul 12;10:36.

21. DiTroia SP, Percharde M, Guerquin M-J, Wall E, Collignon E, Ebata KT, et al. Maternal vitamin C regulates reprogramming of DNA methylation and germline development. Nature. 2019 Sep 4;573(7773):271–5.

22. Yue X, Rao A. TET family dioxygenases and the TET activator vitamin C in immune responses and cancer. Blood. 2020 Sep 17;136(12):1394–401.

23. Gustafson CB, Yang C, Dickson KM, Shao H, Van Booven D, Harbour JW, et al. Epigenetic reprogramming of melanoma cells by vitamin C treatment. Clin Epigenetics. 2015 Apr 29;7(1):51.

24. Agathocleous M, Meacham CE, Burgess RJ, Piskounova E, Zhao Z, Crane GM, et al. Ascorbate regulates haematopoietic stem cell function and leukaemogenesis. Nature. 2017 Sep 28;549(7673):476–81.

25. Bourque G, Burns KH, Gehring M, Gorbunova V, Seluanov A, Hammell M, et al. Ten things you should know about transposable elements. Genome Biol. 2018 Nov 19;19(1):199.

26. Klawitter S, Fuchs NV, Upton KR, Muñoz-Lopez M, Shukla R, Wang J, et al. Reprogramming triggers endogenous L1 and Alu retrotransposition in human induced pluripotent stem cells. Nat Commun. 2016 Jan 8;7:10286.

27. Gerdes P, Lim SM, Ewing AD, Larcombe MR, Chan D, Sanchez-Luque FJ, et al. Retrotransposon instability dominates the acquired mutation landscape of mouse induced pluripotent stem cells. Nat Commun. 2022 Dec 3;13(1):7470.

28. Wissing S, Muñoz-Lopez M, Macia A, Yang Z, Montano M, Collins W, et al. Reprogramming somatic cells into iPS cells activates LINE-1 retroelement mobility. Hum Mol Genet. 2012 Jan 1;21(1):208–18.

29. Fueyo R, Judd J, Feschotte C, Wysocka J. Roles of transposable elements in the regulation of mammalian transcription. Nat Rev Mol Cell Biol. 2022 Jul;23(7):481– 97.

30. Lawson HA, Liang Y, Wang T. Transposable elements in mammalian chromatin organization. Nat Rev Genet. 2023 Jun 7;

31. Gazquez-Gutierrez A, Witteveldt J R Heras S, Macias S. Sensing of transposable elements by the antiviral innate immune system. RNA. 2021 Apr 22;27(7):735–52.

32. de la Rica L, Deniz Ö Cheng KCL, Todd CD, Cruz C, Houseley J, et al. TET-dependent regulation of retrotransposable elements in mouse embryonic stem cells. Genome Biol. 2016 Nov 18;17(1):234.

33. Deniz Ö, de la Rica L, Cheng KCL, Spensberger D, Branco MR. SETDB1 prevents TET2-dependent activation of IAP retroelements in naïve embryonic stem cells. Genome Biol. 2018 Jan 19;19(1):6.

34. Walter M, Teissandier A, Pérez-Palacios R, Bourc’his D. An epigenetic switch ensures transposon repression upon dynamic loss of DNA methylation in embryonic stem cells. Elife. 2016 Jan 27;5.

35. Liu M, Ohtani H, Zhou W, Ørskov AD, Charlet J, Zhang YW, et al. Vitamin C increases viral mimicry induced by 5-aza-2’-deoxycytidine. Proc Natl Acad Sci USA. 2016 Sep 13;113(37):10238–44.

36. Silva J, Barrandon O, Nichols J, Kawaguchi J, Theunissen TW, Smith A. Promotion of reprogramming to ground state pluripotency by signal inhibition. PLoS Biol. 2008 Oct 21;6(10):e253.

37. Carey TS, Cao Z, Choi I, Ganguly A, Wilson CA, Paul S, et al. BRG1 governs nanog transcription in early mouse embryos and embryonic stem cells via antagonism of histone H3 lysine 9/14 acetylation. Mol Cell Biol. 2015 Dec;35(24):4158–69.

38. Yang WR, Ardeljan D, Pacyna CN, Payer LM, Burns KH. SQuIRE reveals locus-specific regulation of interspersed repeat expression. Nucleic Acids Res. 2019 Mar 18;47(5):e27.

39. Gagnier L, Belancio VP, Mager DL. Mouse germ line mutations due to retrotransposon insertions. Mob DNA. 2019 Apr 13;10:15.

40. de la Rica L, Stanley JS, Branco MR. Profiling DNA methylation and hydroxymethylation at retrotransposable elements. Methods Mol Biol. 2016;1400:387–401.

41. Hu X, Zhang L, Mao S-Q, Li Z, Chen J, Zhang R-R, et al. Tet and TDG mediate DNA demethylation essential for mesenchymal-to-epithelial transition in somatic cell reprogramming. Cell Stem Cell. 2014 Apr 3;14(4):512–22.

42. Tsumura A, Hayakawa T, Kumaki Y, Takebayashi S, Sakaue M, Matsuoka C, et al. Maintenance of self-renewal ability of mouse embryonic stem cells in the absence of DNA methyltransferases Dnmt1, Dnmt3a and Dnmt3b. Genes Cells. 2006 Jul;11(7):805–14.

43. Robbez-Masson L, Tie CHC, Conde L, Tunbak H, Husovsky C, Tchasovnikarova IA, et al. The HUSH complex cooperates with TRIM28 to repress young retrotransposons and new genes. Genome Res. 2018 Jun;28(6):836–45.

44. Rowe HM, Jakobsson J, Mesnard D, Rougemont J, Reynard S, Aktas T, et al. KAP1 controls endogenous retroviruses in embryonic stem cells. Nature. 2010 Jan 14;463(7278):237–40.

45. Tomaz RA, Harman JL, Karimlou D, Weavers L, Fritsch L, Bou-Kheir T, et al. Jmjd2c facilitates the assembly of essential enhancer-protein complexes at the onset of embryonic stem cell differentiation. Development. 2017 Feb 15;144(4):567–79.

46. Pedersen MT, Kooistra SM, Radzisheuskaya A, Laugesen A, Johansen JV, Hayward DG, et al. Continual removal of H3K9 promoter methylation by Jmjd2 demethylases is vital for ESC self-renewal and early development. EMBO J. 2016 Jul 15;35(14):1550–64.

47. Iwamori N, Zhao M, Meistrich ML, Matzuk MM. The testis-enriched histone demethylase, KDM4D, regulates methylation of histone H3 lysine 9 during spermatogenesis in the mouse but is dispensable for fertility. Biol Reprod. 2011 Jun;84(6):1225–34.

48. Liu S, Brind’Amour J, Karimi MM, Shirane K, Bogutz A, Lefebvre L, et al. Setdb1 is required for germline development and silencing of H3K9me3-marked endogenous retroviruses in primordial germ cells. Genes Dev. 2014 Sep 15;28(18):2041–55.

49. Matsui T, Leung D, Miyashita H, Maksakova IA, Miyachi H, Kimura H, et al. Proviral silencing in embryonic stem cells requires the histone methyltransferase ESET. Nature. 2010 Apr 8;464(7290):927–31.

50. Karimi MM, Goyal P, Maksakova IA, Bilenky M, Leung D, Tang JX, et al. DNA methylation and SETDB1/H3K9me3 regulate predominantly distinct sets of genes, retroelements, and chimeric transcripts in mESCs. Cell Stem Cell. 2011 Jun 3;8(6):676–87.

51. Streva VA, Jordan VE, Linker S, Hedges DJ, Batzer MA, Deininger PL. Sequencing, identification and mapping of primed L1 elements (SIMPLE) reveals significant variation in full length L1 elements between individuals. BMC Genomics. 2015 Mar 21;16(1):220.

52. Witherspoon DJ, Xing J, Zhang Y, Watkins WS, Batzer MA, Jorde LB. Mobile element scanning (ME-Scan) by targeted high-throughput sequencing. BMC Genomics. 2010 Jun 30;11:410.

53. Navarro FC, Hoops J, Bellfy L, Cerveira E, Zhu Q, Zhang C, et al. TeXP: Deconvolving the effects of pervasive and autonomous transcription of transposable elements. PLoS Comput Biol. 2019 Aug 19;15(8):e1007293.

54. Hajkova P. Epigenetic reprogramming--taking a lesson from the embryo. Curr Opin Cell Biol. 2010 Jun;22(3):342–50.

55. Takahashi K, Tanabe K, Ohnuki M, Narita M, Ichisaka T, Tomoda K, et al. Induction of pluripotent stem cells from adult human fibroblasts by defined factors. Cell. 2007 Nov 30;131(5):861–72.

56. Takahashi K, Yamanaka S. Induction of pluripotent stem cells from mouse embryonic and adult fibroblast cultures by defined factors. Cell. 2006 Aug 25;126(4):663–76.

57. Ying Q-L, Wray J, Nichols J, Batlle-Morera L, Doble B, Woodgett J, et al. The ground state of embryonic stem cell self-renewal. Nature. 2008 May 22;453(7194):519–23.

58. Castro-Diaz N, Ecco G, Coluccio A, Kapopoulou A, Yazdanpanah B, Friedli M, et al. Evolutionally dynamic L1 regulation in embryonic stem cells. Genes Dev. 2014 Jul 1;28(13):1397–409.

59. Sookdeo A, Hepp CM, McClure MA, Boissinot S. Revisiting the evolution of mouse LINE-1 in the genomic era. Mob DNA. 2013 Jan 3;4(1):3.

60. Naas TP, DeBerardinis RJ, Moran JV, Ostertag EM, Kingsmore SF, Seldin MF, et al. An actively retrotransposing, novel subfamily of mouse L1 elements. EMBO J. 1998 Jan 15;17(2):590–7.

61. Bulut-Karslioglu A, De La Rosa-Velázquez IA, Ramirez F, Barenboim M, Onishi-Seebacher M, Arand J, et al. Suv39h-dependent H3K9me3 marks intact retrotransposons and silences LINE elements in mouse embryonic stem cells. Mol Cell. 2014 Jul 17;55(2):277–90.

62. Percharde M, Lin C-J, Yin Y, Guan J, Peixoto GA, Bulut-Karslioglu A, et al. A LINE1-Nucleolin Partnership Regulates Early Development and ESC Identity. Cell. 2018 Jul 12;174(2):391–405.e19.

63. Love MI, Huber W, Anders S. Moderated estimation of fold change and dispersion for RNA-seq data with DESeq2. Genome Biol. 2014;15(12):550.

64. Jin Y, Tam OH, Paniagua E, Hammell M. TEtranscripts: a package for including transposable elements in differential expression analysis of RNA-seq datasets. Bioinformatics. 2015 Nov 15;31(22):3593–9.

65. Langmead B, Salzberg SL. Fast gapped-read alignment with Bowtie 2. Nat Methods. 2012 Mar 4;9(4):357–9.

66. Zhang Y, Liu T, Meyer CA, Eeckhoute J, Johnson DS, Bernstein BE, et al. Model-based analysis of ChIP-Seq (MACS). Genome Biol. 2008 Sep 17;9(9):R137.

67. Krueger F, Andrews SR. Bismark: a flexible aligner and methylation caller for Bisulfite-Seq applications. Bioinformatics. 2011 Jun 1;27(11):1571–2.

