## Additional File 1 for "Vitamin C activates young LINE-1 elements in mouse embryonic stem cells via H3K9me3 demethylation"

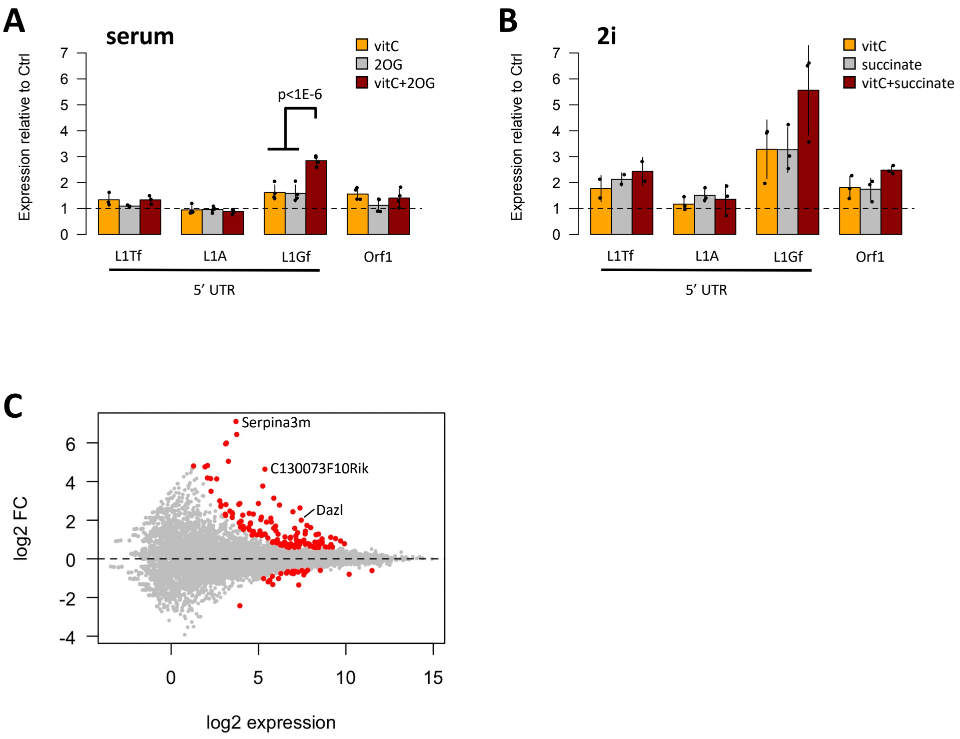


**Figure S1.** A) qRT-PCR data for L1 expression analysis following addition of 2OG to serum media with or without vitC. P-value is from an ANOVA with Tukey's post-hoc test. B) qRT-PCR data for L1 expression in 2i-grown ESC following succinate addition with or without vitC. ANOVA with Tukey's post-hoc test gave no p-values below 0.05. C) MA plot for gene expression from RNA-seq of vitC-treated 2i-grown cells. Differentially expressed genes are highlighted in red.

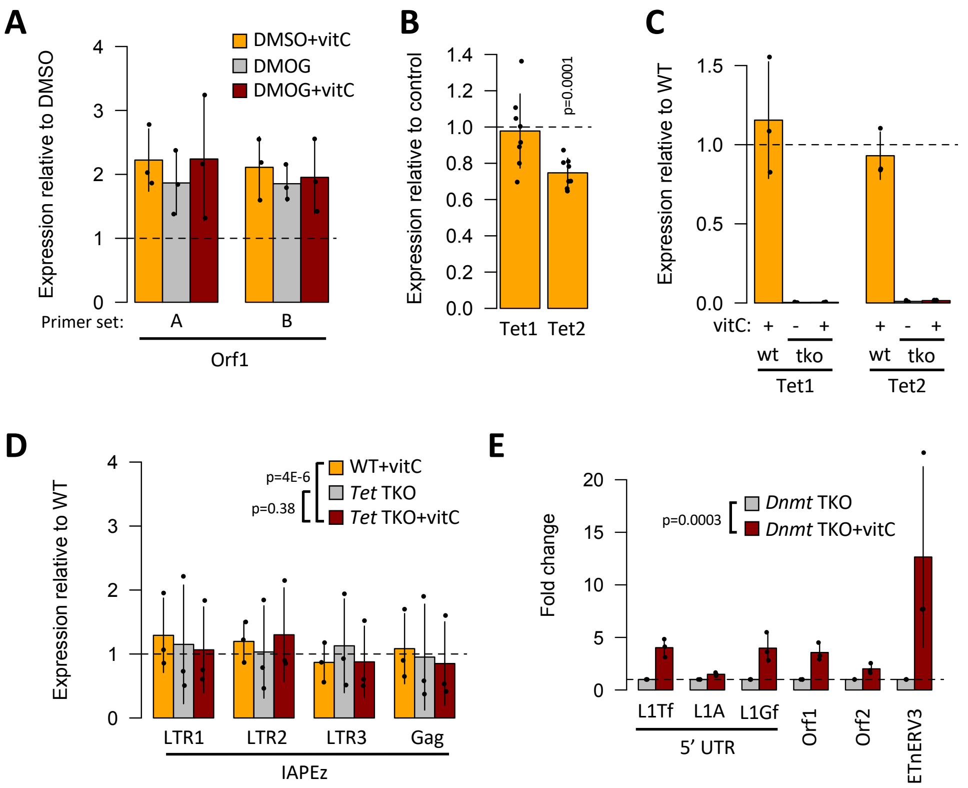


**Figure S2.** A) qRT-PCR data for L1 Orf1 expression in DMOG-treated cells compared to DMSO controls, with and without vitC. ANOVA revealed no significant differences in L1 upregulation between the groups. B) qRT-PCR data showing that *Tet1/2* transcripts did not increase following vitC treatment C) qRT-PCR data (normalised to untreated WT cells) confirming lack of *Tet1* and *Tet2* expression in *Tet* triple knock-out (TKO) mESCs. D) qRT-PCR data for IAP elements in WT and TKO Tet TKO cells. P-values are from an ANOVA model. E) qRT-PCR data showing L1 expression in *Dnmt1* TKO cells. P-value is from an ANOVA model.

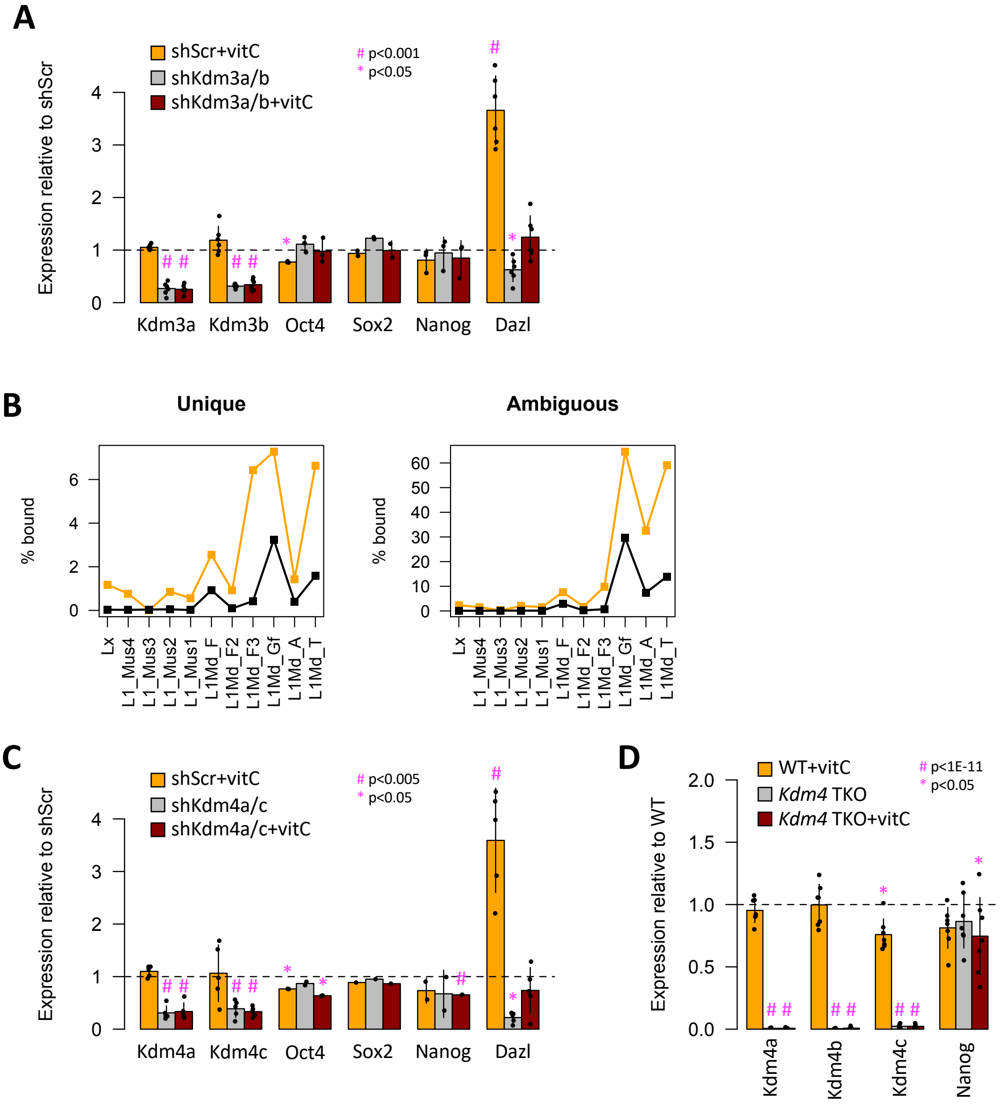


**Figure S3.** A) qRT-PCR data for *Kdm3a/b*, pluripotency markers and *Dazl* (positive control) in *Kdm3a/b* KD mESCs. B) ChIP-seq data for KDM4A (Pedersen et al., 2016) and KDM4C (Tomaz et al., 2017) showing the % of bound protein at different L1 families in the mouse genome, for peaks from both uniquely (left chart) or ambiguously (right chart) aligned reads. C) qRT-PCR data for *Kdm4a/c*, pluripotency markers and *Dazl* (positive control) in *Kdm4a/c* KD mESCs. D) qRT-PCR data showing expression of *Kdm4a/b/c* and *Nanog* in inducible *Kdm4a/b/c* triple KO (TKO) mESCs. All p-values shown are from one-sample t-tests (mu=1) with Benjamini-Hochberg multiple comparisons correction.

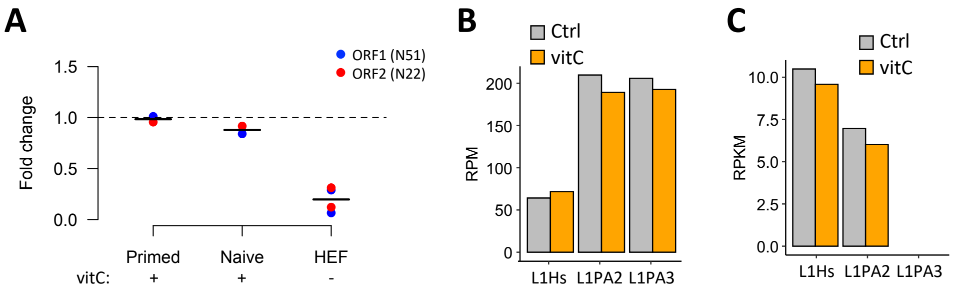


**Figure S4.** A) qRT-PCR data for the same samples shown in Figure 6C (ORF1p western blot), normalised to untreated hESCs. Expression data from human fibroblasts (HEF) are shown for comparison. B) Analysis of RNA-seq data in control and 24h vitC-treated primed hESCs by TEtranscripts, showing subfamily level expression of young L1s. C) As in B, but using TeXP for analysis.
