## Additional File 5 for "Vitamin C activates young LINE-1 elements in mouse embryonic stem cells via H3K9me3 demethylation"

### Sequencing strategy for detection of mouse L1 insertions

### The sequencing approach consisted of a library preparation including a single-step PCR amplification and enrichment of junctions of the different L1 families present in the mouse genome. In general, the DNA shearing was done through sonication and the library was prepared with the reagents provided by the Illumina TruSeq DNA LT Nano library prep kit. However, homemade adapters were prepared in order to incorporate only one of the apex sequences necessary for cluster generation, so the second apex sequence is introduced as a 5’-flap in primers binding both the 5’ and 3’ L1 termini sequences in outward directions. Thus, the only fragments that would be amplified during the library prep PCR will be those both containing the junction of a L1 copy and successfully bound to an adapter, leading to a simultaneous enrichment in L1 junctions. Additionally, since the Illumina index is usually contained in the apex sequence eliminated from our adapters, the indices were incorporated at the end of the ‘read 1’ sequence of the adapter and they will be read within the read 1 sequencing, immediately before the sequence of each fragment. Thus, the adapters were generated by annealing of two primers obtained from IDT with the following structures:

### AATGATACGGCGACCACCGAGATCTACACTCTTTCCCTACACGACGCTCTTCCGATCT[6ntBCs](C)*T

### [5Phos](G)[6ntBCas]AGATCGGAAGAGCGTCGTGTAGGGAAAGAGTGTAGA[SpcC3]

### The asterisk represents a phosphorothioate linkage, [5Phos] indicates a 5’ phosphate end, [6ntBCs] indicates the sense sequence of each of the 24 6nt-long barcodes of the low throughput (LT) Illumina TruSeq kits (for instance, barcode 6 sequence is GCCAAT) and the [6ntBCas] represents the complementary sequence of the corresponding barcode. The (C) and (G) are an extra nucleotide that was added to half of the adapters in order to create diversity in the 7th position of the read 1, and was differentially included in barcode pairs that are recommended to be pooled together by Illumina guidelines (for instance, adapter with barcode 6 had the extra nucleotide and adapter with barcode 12 did not). The functional adapters were obtained by combining the primer pairs at a final concentration of 15µM each, denaturing them by incubation at 95ºC for 2 min and letting the block to cool down to room temperature (~25ºC).

The sequencing run was configured for using custom index read primer and custom read 2 primers following manufacturer’s instructions. The custom index read primer (ACACTCTTTCCCTACACGACGCTCTTCCGATCT) was loaded in the kit’s cartridge using manufacturer’s indications. Note that this primer is the same as the standard Illumina read 1 primer in order to re-read the barcodes that are included at the beginning of the read 1 sequence and use the instrument software to sort the barcoded sequences. The custom read 2 primer was a mix of MouseRead2_3end primer, Adapter_mL1Tf_5end, Adapter_mL1Tf3_5end, Adapter_mL1Gf_5end, Adapter_mL1TGf_5end2, Adapter_mL1A_5end1 and Adapter_mL1A_5end2 (Supplementary Table 4) at a ratio 6:1:1:1:1:1:1, using the total concentration of the pool as reference for loading the mix in the kit cartridge using manufacturer’s indications.
